## Supplementary material for "Genome graphs reveal the importance of structural variation in *Mycobacterium tuberculosis* evolution and drug resistance": Fig.S

#### Contents


- Supplementary Figures
- Supplementary Tables
- Supplementary Results
- Supplementary Methods
- Supplementary References

#### Supplementary Figures

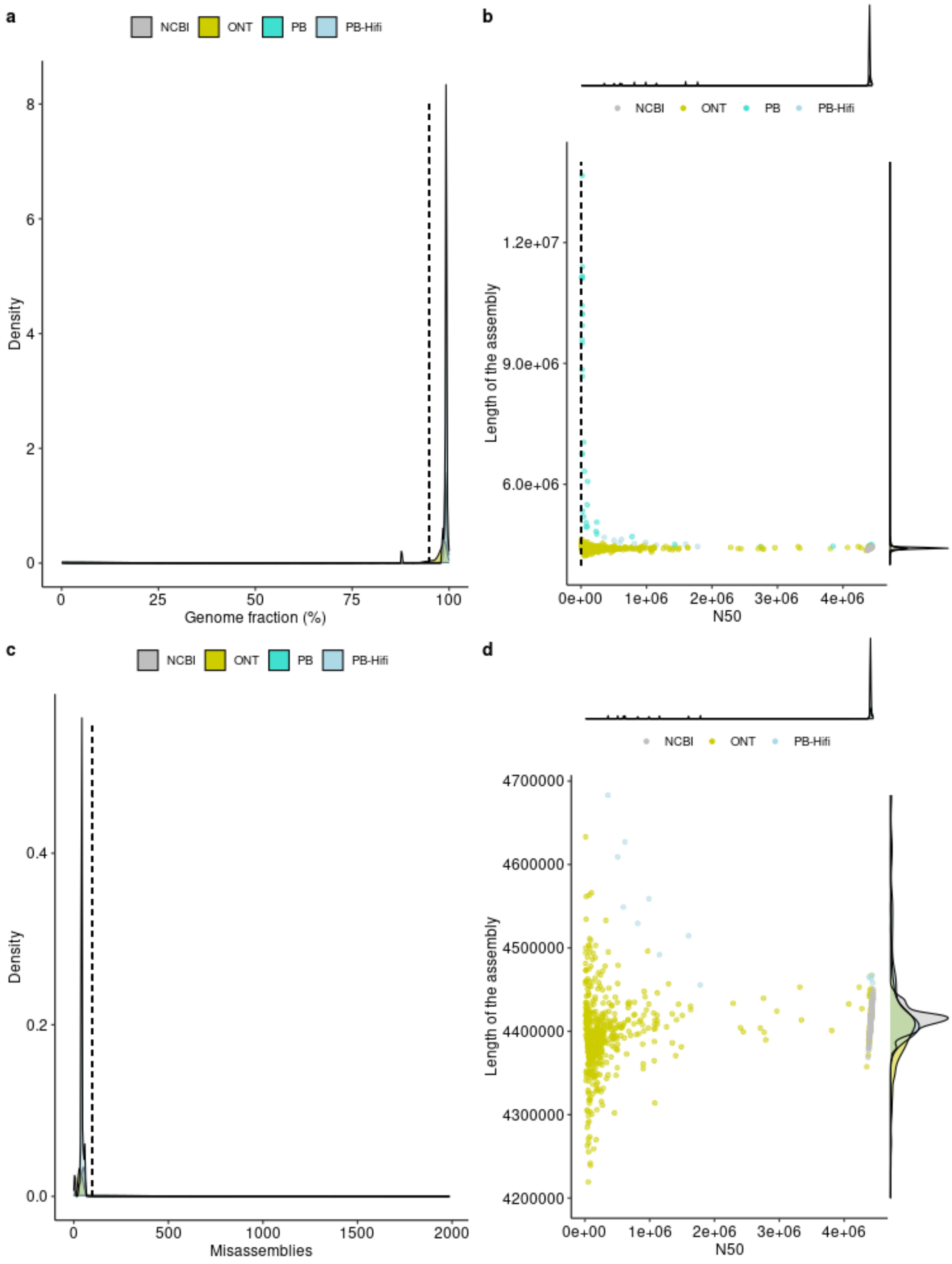


**Supplementary Figure 1.** QUAST metrics used to filter out assemblies. **a**, A density plot showing the trends of genome fraction as a percentage grouped by the genome technology used to obtain it (or whether it is an assembly downloaded directly from NCBI); the dashed line represents the 95% threshold used to filter out samples. **b**, A scatter plot where the N50 of each value is on the x axis while the total assembly length is on the y axis grouped by the genome technology; the dashed line represents the 10,000bp N50 threshold used to filter out samples. On the edge of each axis the density plots are displayed for both the N50 and the assembly length to further study the different distributions across genome technologies. **c**, A density plot shows the distribution of misassemblies across all assemblies, grouped by the genome technology; the dashed line represent the 100 number of misassemblies filter applied to this data. **d**, A similar plot to b with the final, filtered data.

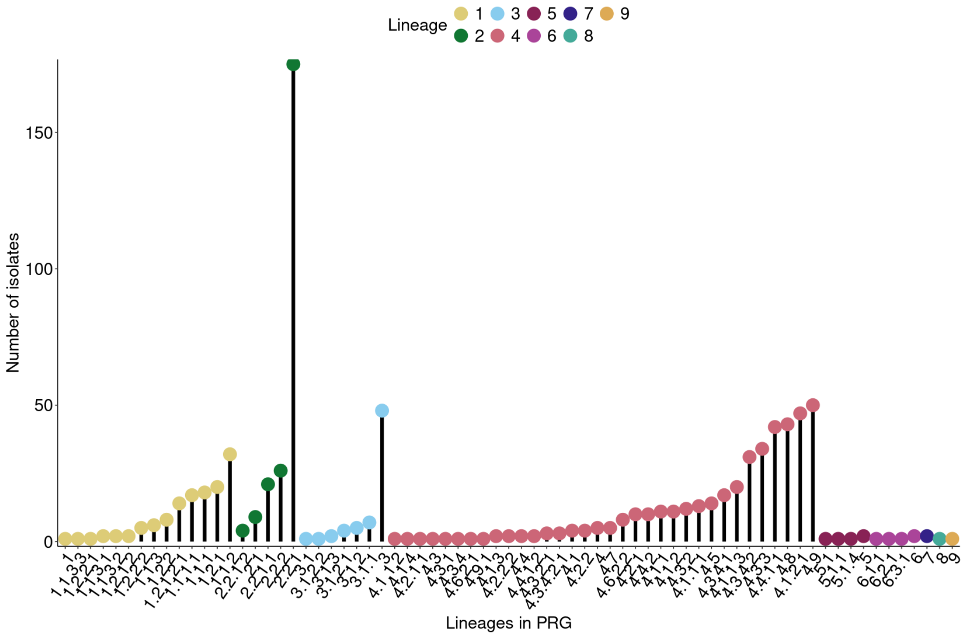


**Supplementary Figure 2.** Lollipop plot of lineage frequency in final list of isolates.

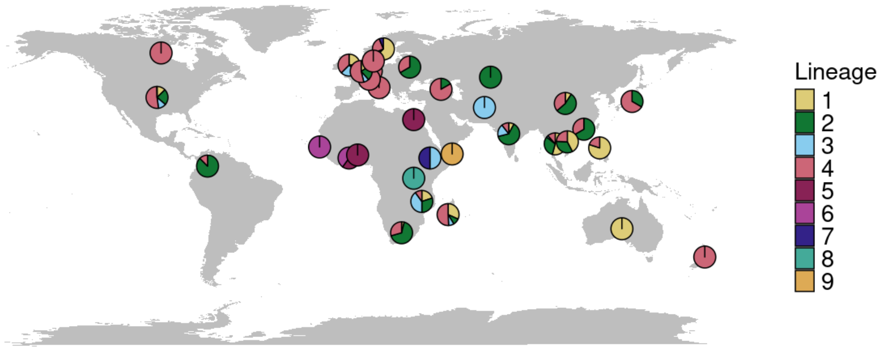


**Supplementary Figure 3.** Map of the origin of the samples (where applicable) or the sequencing location. Pie charts represent the frequency of lineages observed per country.

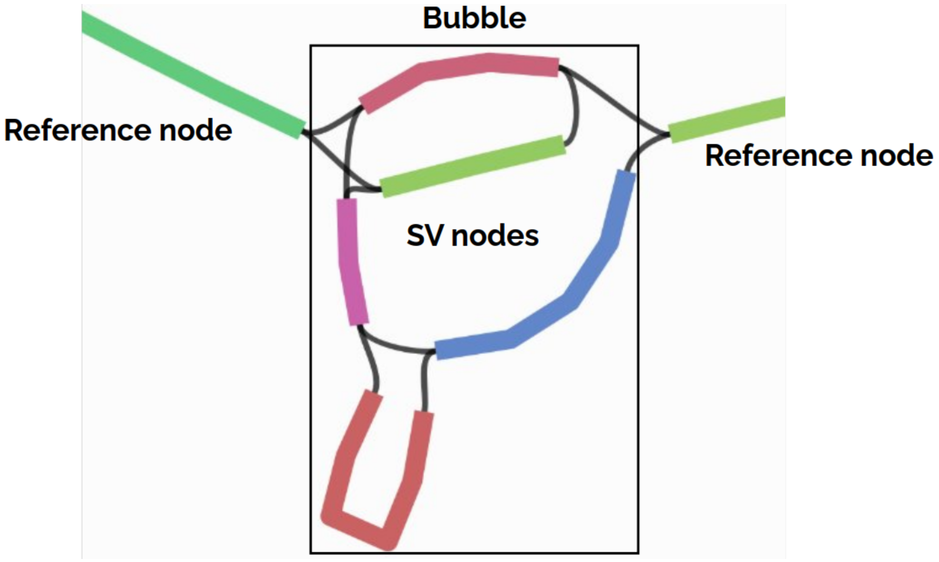


**Supplementary Figure 4.** Visualization of nodes forming a bubble in the PRG. A node is the terminology used for any sequence present in the graph; a reference node is a sequence that is conserved across all isolates while an SV node is a sequence that is variable and not found across all isolates. A group of nodes that are connected to each other and are not conserved across all isolates form a bubble. Visualization done using Bandage [17].

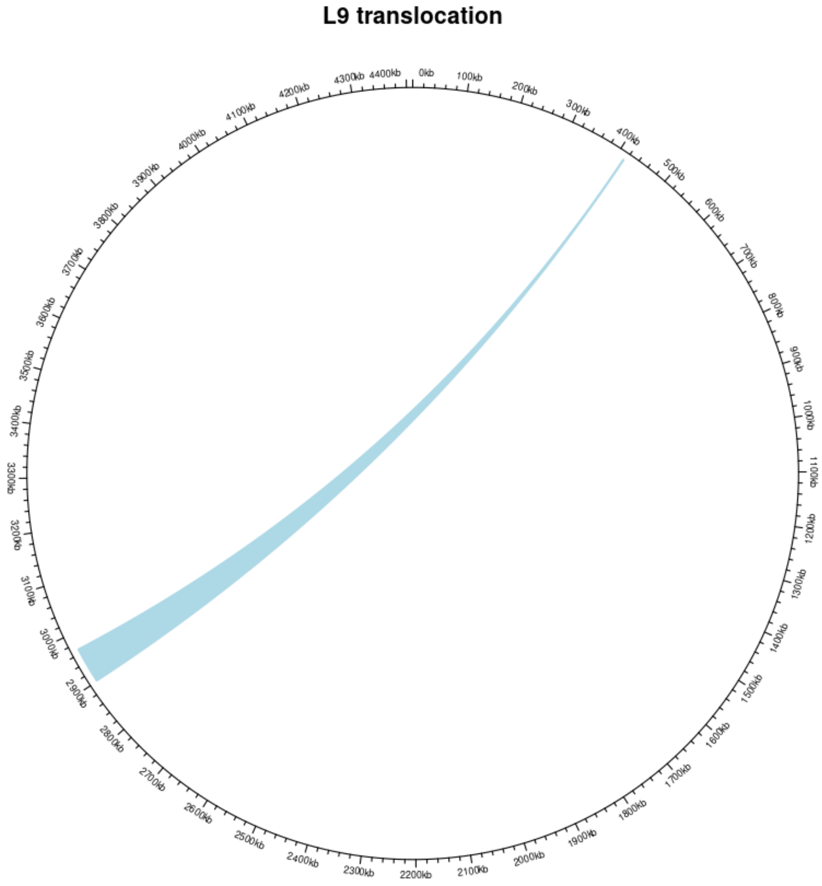


**Supplementary Figure 5.** Translocation of the 66kbp region in the L9 genome.

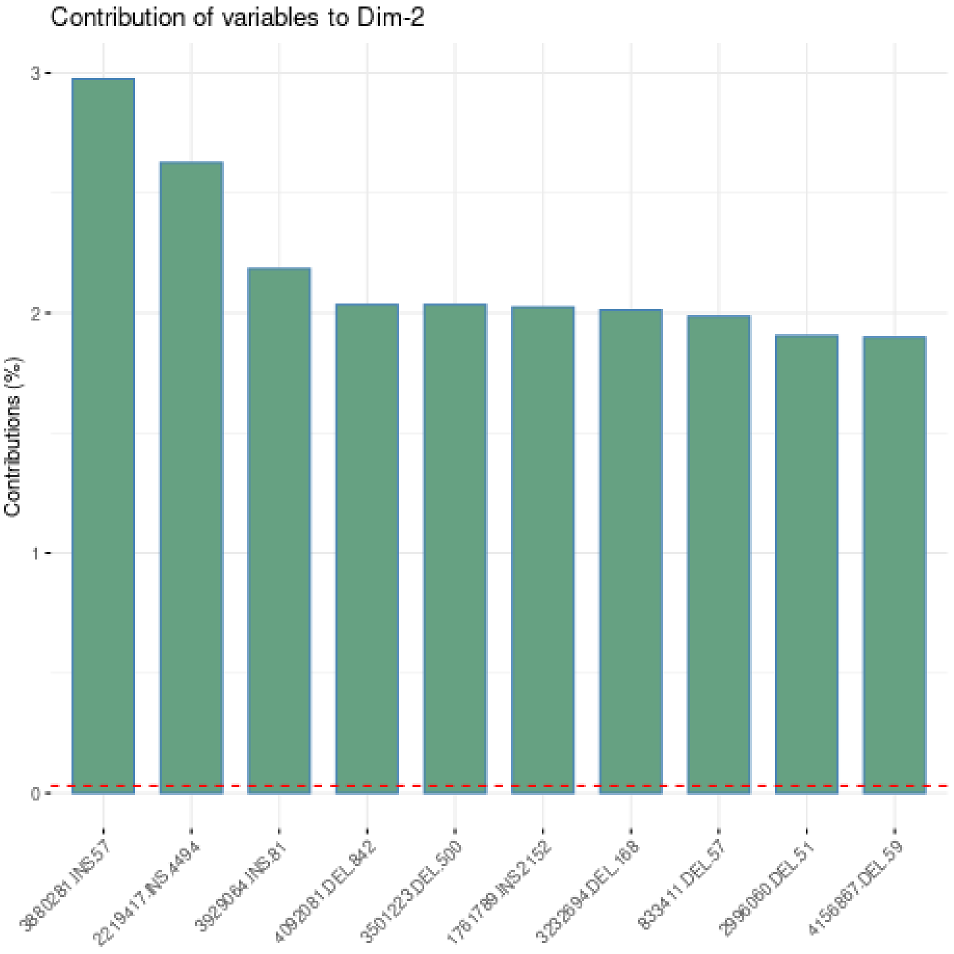


**Supplementary Figure 6.** Top 10 SVs that contribute to the variability of principal component 2 (PC2) from the principal component analysis (PCA) done using the *Mtb* SV genotypes. 1761789.INS.2152 represents the TbD1 SV. The reference, red, dashed line corresponds to the expected value if the contribution where uniform.

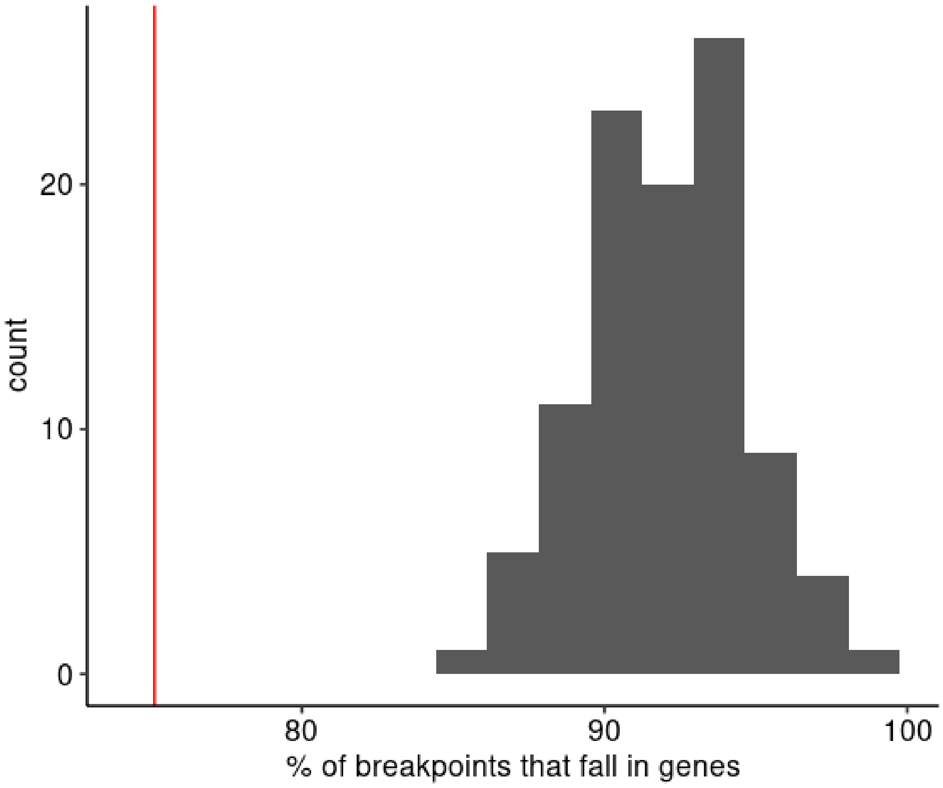


**Supplementary Figure 7.** Histogram of percentage of simulated random breakpoints that fall in genes in the *Mtb* genome out of 100, permuted for 100 iterations. Vertical red line represents the observed percentage of SV breakpoints that fall in genes.

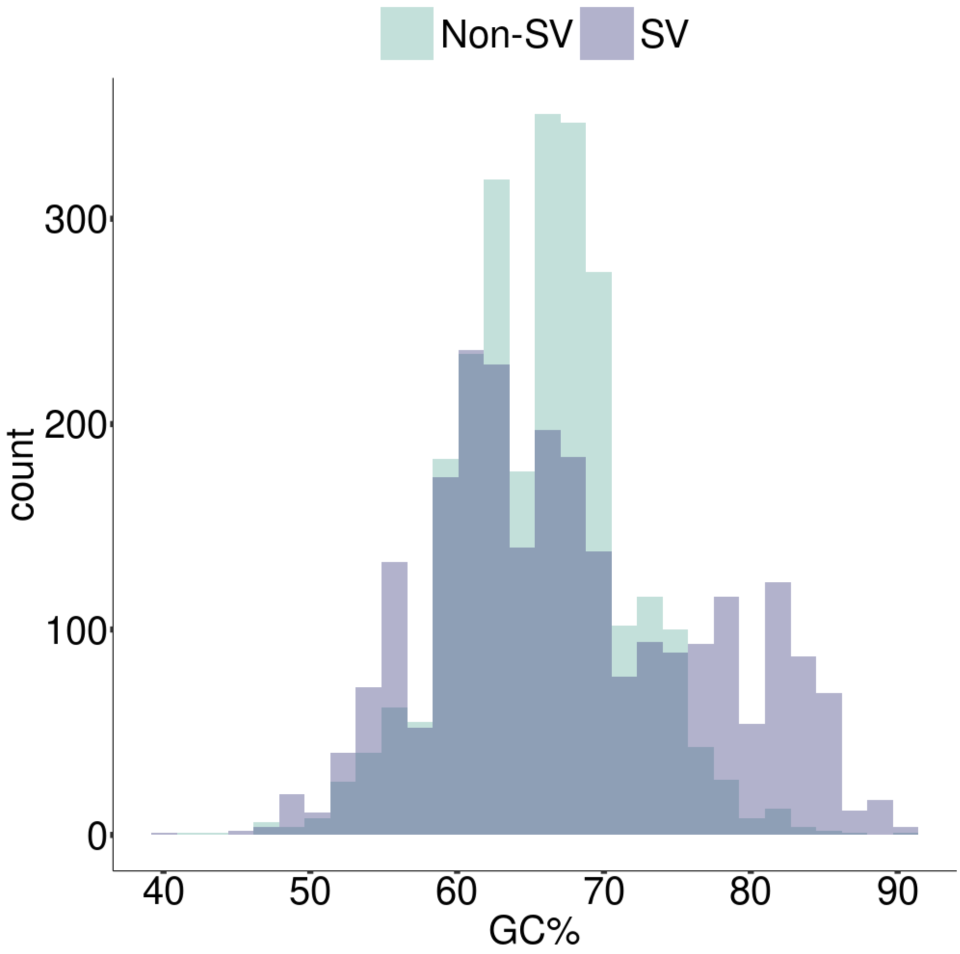


**Supplementary Figure 8.** Histogram depicting the GC% of the SV breakpoints compared to randomly selected 100-bp regions from the *Mtb* genome.

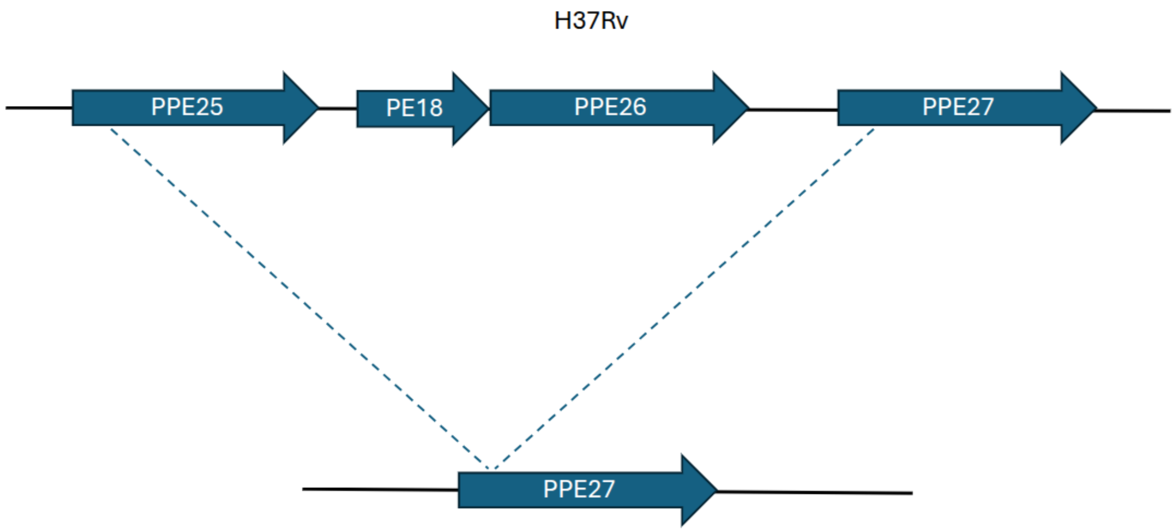


**Supplementary Figure 9.** Schematic of the *ppe25*-*ppe27* deletion as compared to the H37Rv genome. The start of both *PPE25* and *PPE27* is homologous, so a deletion of the start of *PPE27* can result in a chimeric gene[7].

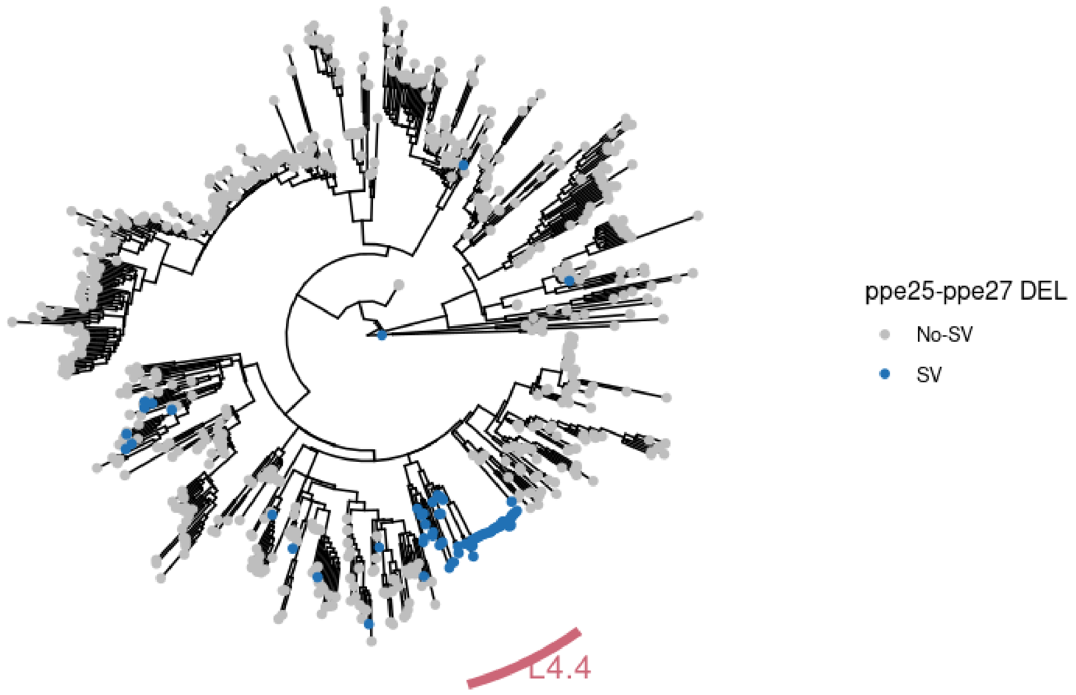


**Supplementary Figure 10.** Mapping the *ppe25*-*ppe27* deletion on the phylogeny created using the SNPs of the *Mtb* long-read genomes included in the *Mtb*-PRG. Colored isolates present the deletion and span lineages 1, 3, 4.1, 4.3, 4.5 and 8.

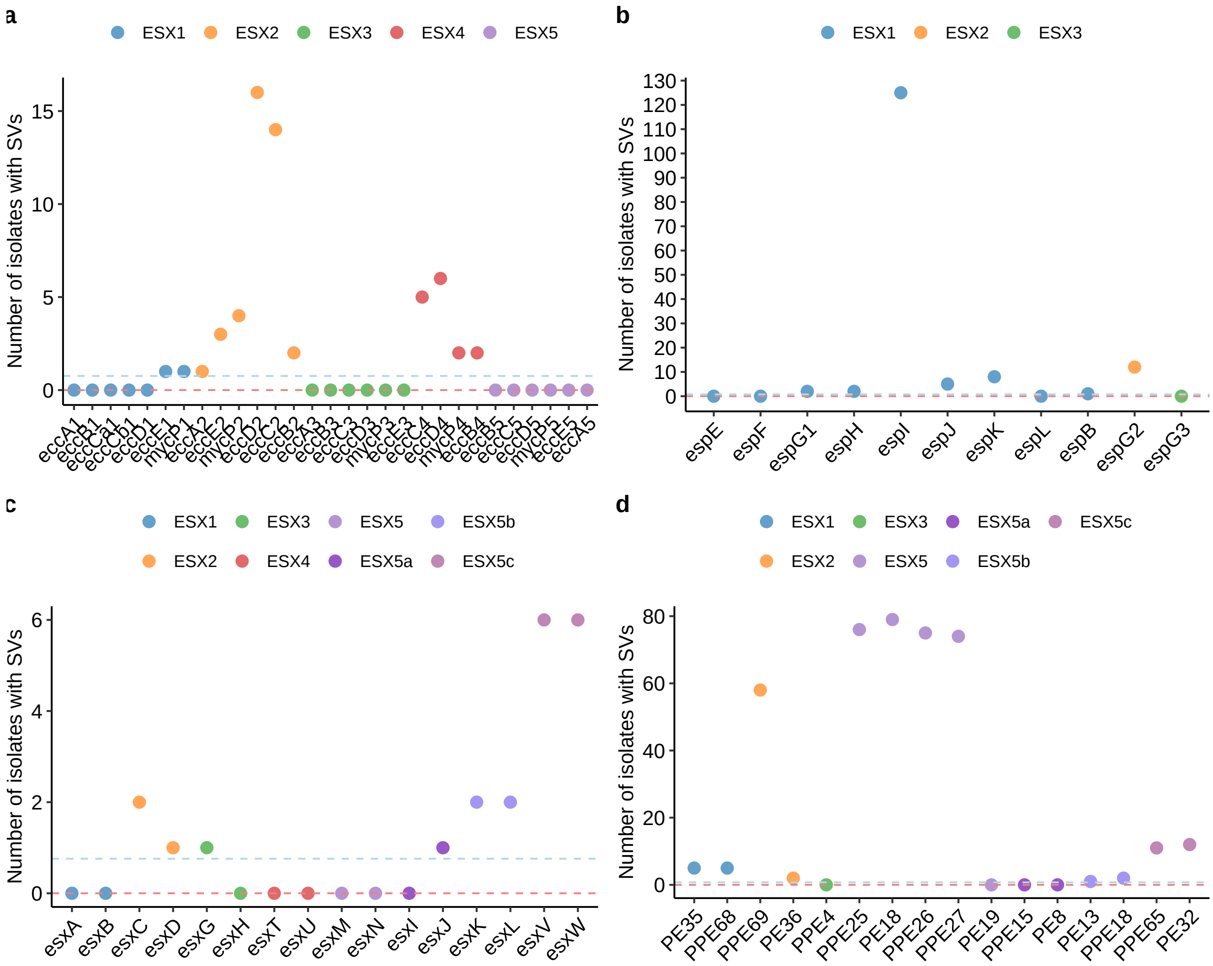


**Supplementary Figure 11.** Number of isolates with SVs across the ESX loci genes.**a-d** Number of isolates with an SV in each gene across the ESX1-5c loci, coloured by the ESX system, grouped into paralogous genes. The red horizontal bar represents the trimmed mean of number of SVs across essential genes; the blue horizontal bar represents the trimmed mean of number of SVs across non-essential genes.

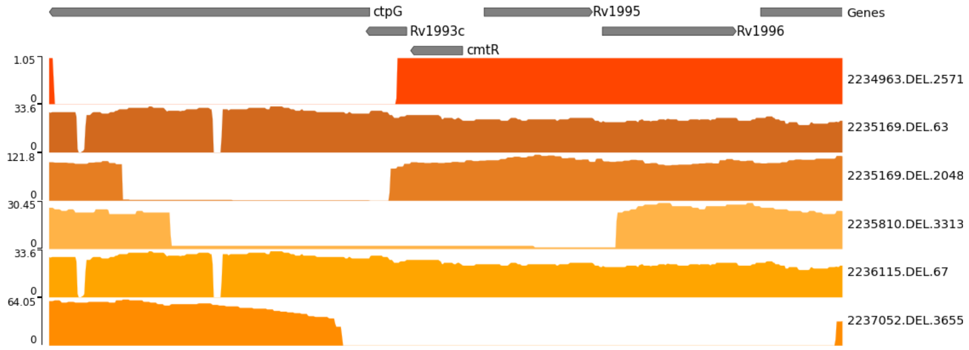


**Supplementary Figure 12.** Genomic long-read coverage of isolates with *ctpG* deletions. Y axis shows the read depth at each genomic position.

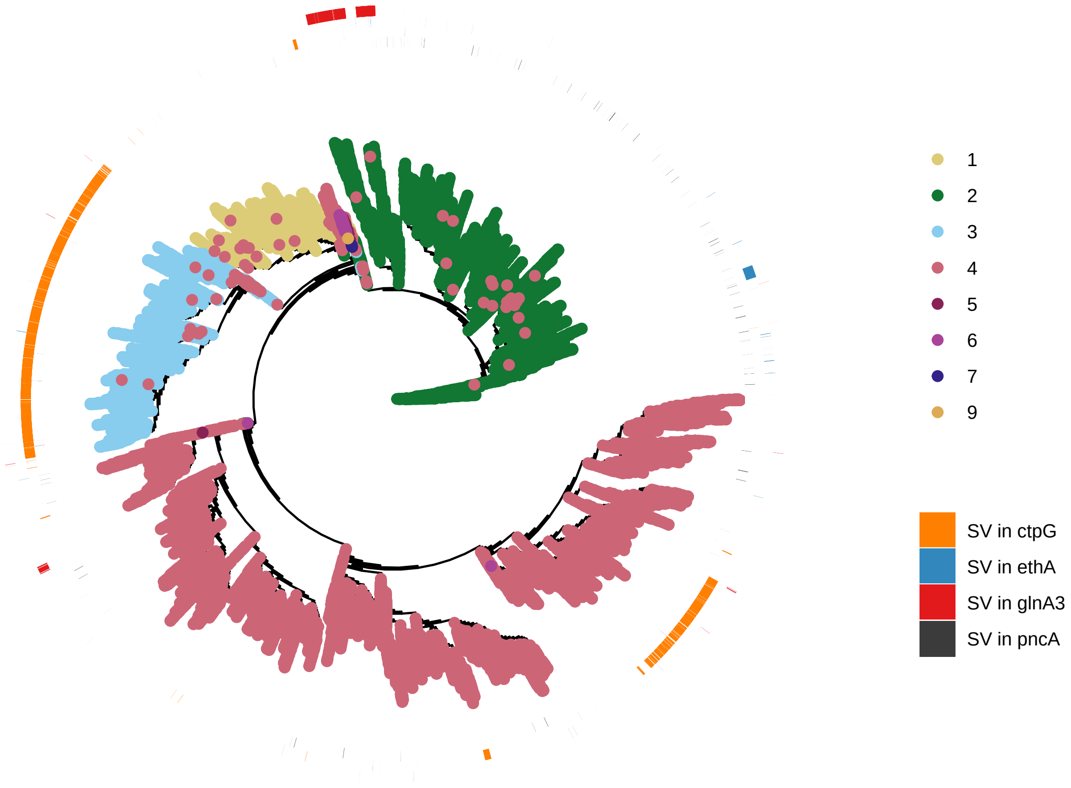


**Supplementary Figure 13.** Homoplasic SVs across 41,134 Illumina-sequenced isolates in an unrooted phylogeny created using their SNPs. The outer rings represent the presence (colour) or absence (white) of an SV in a specific gene for each isolate.

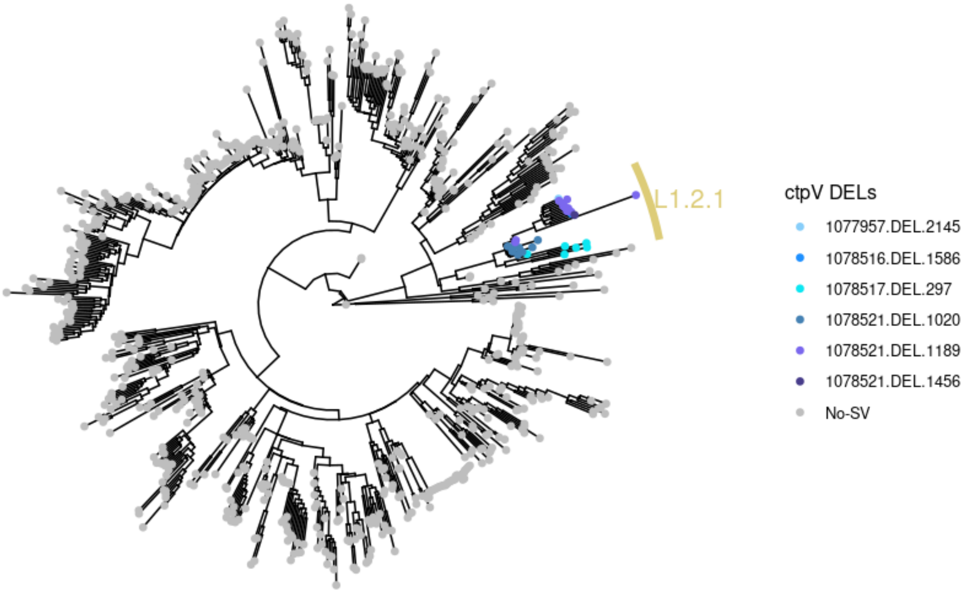


**Supplementary Figure 14.** Mapping the *ctpV* deletions on the phylogeny created using the SNPs of the *Mtb* long-read genomes included in the *Mtb*-PRG.

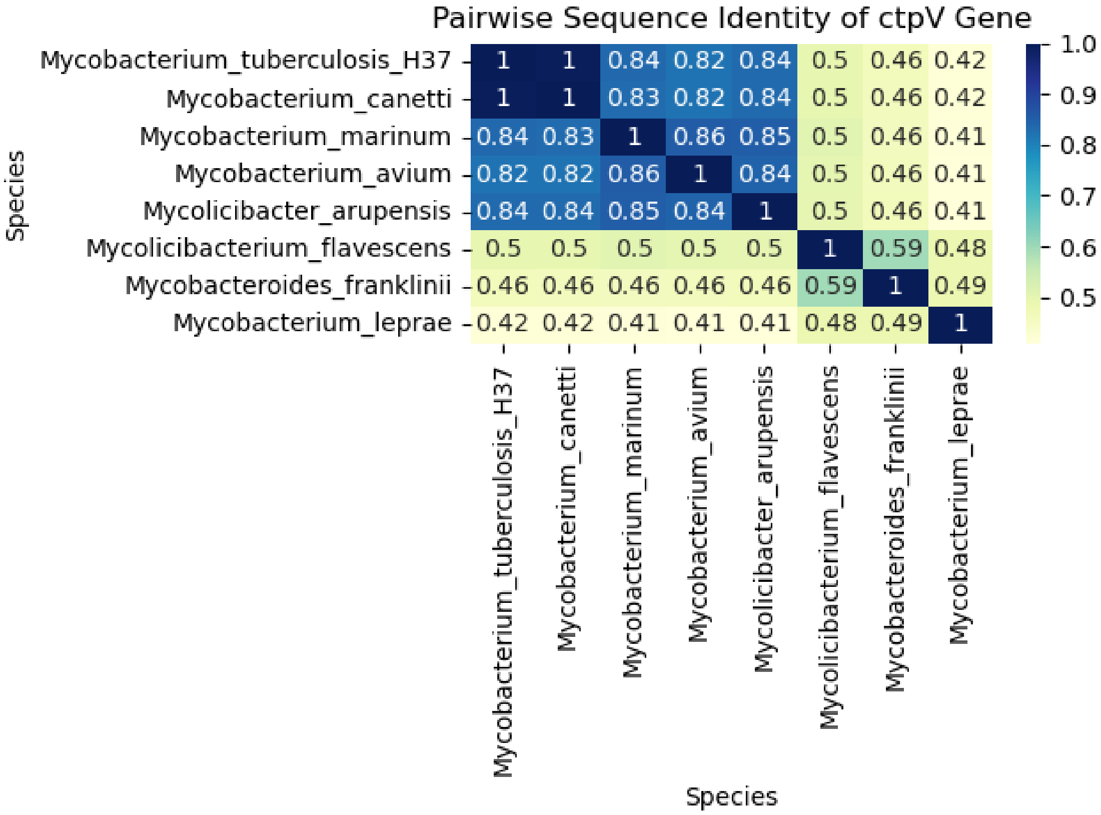


**Supplementary Figure 15.** Pairwise identity of the *ctpV* gene across diverse Mycobacteria.

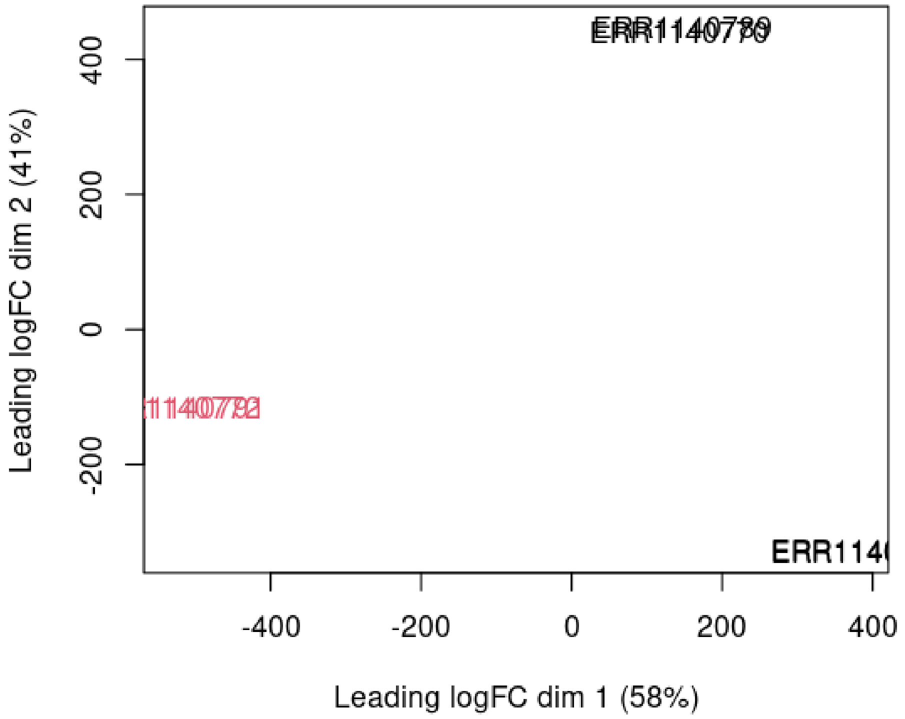


**Supplementary Figure 16.** A multidimensional scaling (MDS) plot. Red-coloured names are L1.2.1. An MDS plot is a visualisation of a principle components analysis (PCA), which determines the greatest sources of variation in the data. This plot shows non-L1.2.1 isolates' transcriptional profiles clustering together, differing from L1.2.1.

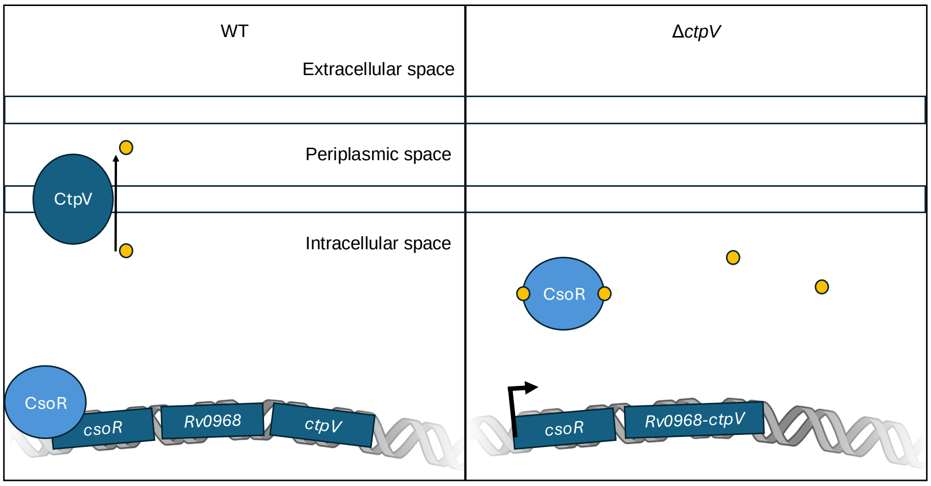


**Supplementary Figure 17.** Proposed hypothetical model explaining the higher expression of *Rv0968* and *ctpV* in L1.2.1 isolates. Under normal conditions, with a functional *ctpV*, the isolate is able to expel the Cu ions, so CsoR inhibits its operon's gene expression (left). In the L1.2.1 isolates, the lack of Cu export leads to its accumulation where CsoR recognizes it, is detached from the DNA and the operon is expressed (right). Yellow circles represent Cu.

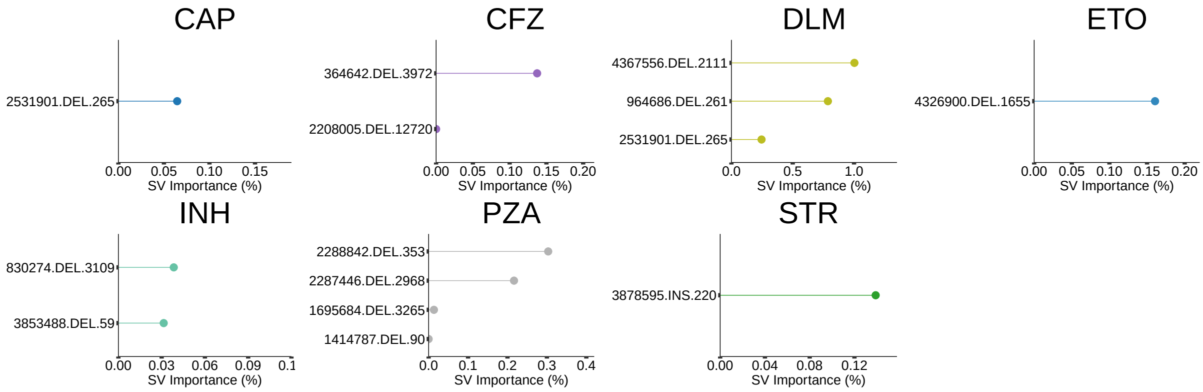


**Supplementary Figure 18.** Machine learning importance scores (%) of significantly associated SVs by drug. Drugs not illustrated did not have any significantly associated SVs. The ID for the SVs is the following: {SV start position in the genome}.{SV type}.{SV size}. CAP: capreomycin, CFZ: clofazimine, DLM: delamanid, ETO: ethionamide, INH: isoniazid, PZA: pyrazinamide, STR: streptomycin.

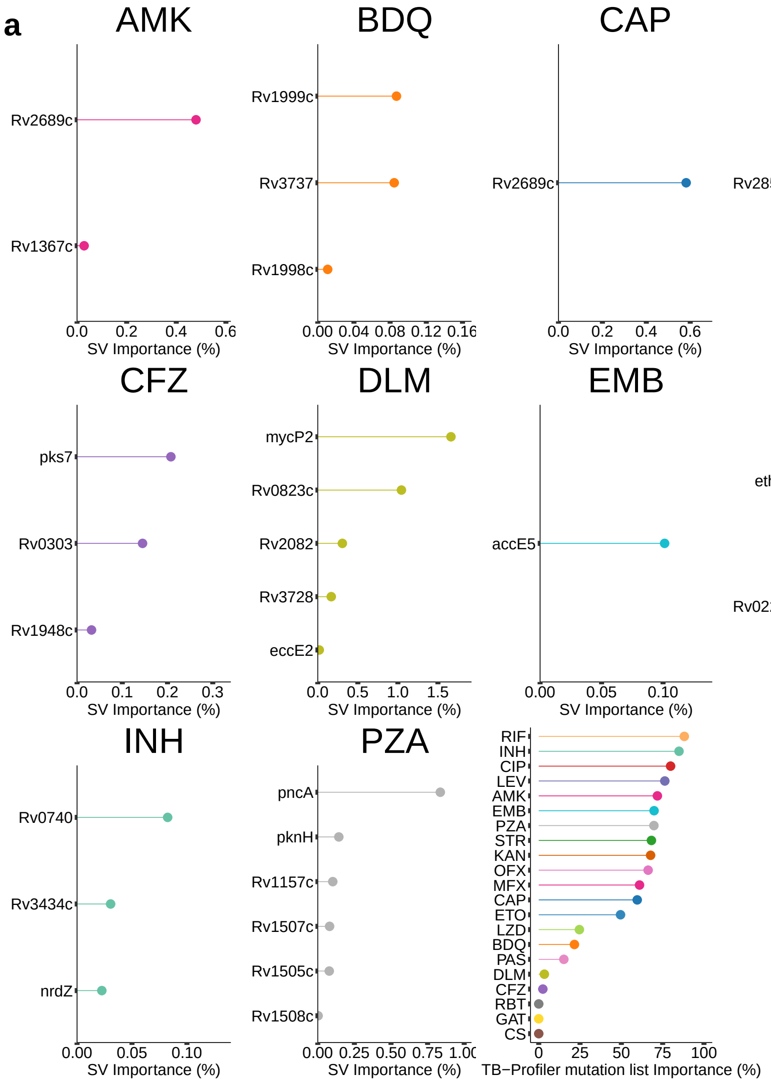


**b**

| Drug | Gene | Start Position | Resistant Frequency | Susceptible Frequency | Total Resistant | Total Susceptible |
|---|---|---|---|---|---|---|
| ETO | Rv0221 | 264067 | 0.13115370 | 0.0239786856 | 2661 | 10134 |
| PZA | pncA | 2288681 | 0.02330152 | 0.0005929440 | 3562 | 16865 |
| INH | Rv3434c | 3853215 | 0.08898369 | 0.0004612740 | 13362 | 23847 |
| PZA | Rv1505c | 1695281 | 0.05434449 | 0.0045363278 | 3257 | 13447 |
| PZA | Rv1508c | 1698095 | 0.05403746 | 0.0038670335 | 3257 | 13447 |
| PZA | Rv1507c | 1696727 | 0.05403746 | 0.0041644977 | 3257 | 13447 |
| AMK | Rv2689c | 3005845 | 0.12210983 | 0.0598475137 | 1384 | 14821 |
| ETO | ethA | 4326004 | 0.05326332 | 0.0034509959 | 2666 | 10142 |
| CAP | Rv2689c | 3005845 | 0.15107103 | 0.0702500253 | 887 | 9879 |
| INH | Rv0740 | 831776 | 0.06587792 | 0.0010179671 | 11157 | 19647 |
| DLM | Rv0823c | 916477 | 0.27102804 | 0.0756793478 | 107 | 7360 |
| DLM | Rv2082 | 2338709 | 0.30769231 | 0.0672163470 | 65 | 5579 |
| PZA | Rv1157c | 1283056 | 0.02020202 | 0.0052271554 | 3168 | 15496 |
| CFZ | Rv0303 | 365234 | 0.01379310 | 0.0006328446 | 145 | 9481 |
| INH | nrdZ | 661295 | 0.01125561 | 0.0045466134 | 13149 | 23534 |
| EMB | accE5 | 3663689 | 0.61379440 | 0.2351801627 | 5321 | 23229 |
| DLM | mycP2 | 4368518 | 0.11290323 | 0.0279764743 | 62 | 6291 |
| DLM | Rv3728 | 4174873 | 0.01869159 | 0.0006771398 | 107 | 7384 |
| DLM | eccE2 | 4366908 | 0.11290323 | 0.0276585598 | 62 | 6291 |
| AMK | Rv1367c | 1539512 | 0.01654846 | 0.0034495503 | 423 | 8117 |

*(preceding page)*: **Supplementary Figure 19.** Genes with resistance-conferring SVs across 41,134 isolates and 22 drugs. **a** Machine learning importance scores (%) of significantly associated genes by drug. Drugs not illustrated did not have any significantly associated genes. **b** Thorough description of the top 20 associated genes, grouped by drugs and ranked by their adjusted p-value. "Total Resistant" is the number of isolates that had a gene phenotype (not an NA) and, therefore, were used in that association test; the same applies to "Total Susceptible". AMK: amikacin, BDQ: bedaquiline, CAP: capreomycin, CFZ: clofazimine, DLM: delamanid, EMB: ethambutol, ETO: ethionamide, INH: isoniazid, LZD: linezolid, OFX: oxfloxacin, PZA: pyrazinamide.

### Supplementary Tables

**Supplementary Table 1.** Lineages of the isolates newly ONT sequenced.

| Lineage | Amount |
|---|---|
| L1.1.1 | 2 |
| L1.1.1.1 | 6 |
| L2.1 | 4 |
| L4.4.2 | 1 |
| L4.5 | 2 |
| L4.6.2.2 | 2 |
| L4.7 | 1 |
| L4.8 | 2 |
| L5 | 2 |
| L6 | 2 |
| L9 | 1 |

**Supplementary Table 2.** Isolates used for the *ctpV* RNA-Seq analysis.

| ID | Lineage | Sequencer | NCBI ID |
|---|---|---|---|
| N0153 | L1.1.1 | Illumina HiSeq 2500 | ERR1140789 |
| N0153 | L1.1.1 | Illumina HiSeq 2500 | ERR1140770 |
| N0072 | L1.1.2 | Illumina HiSeq 2500 | ERR1140766 |
| N0072 | L1.1.2 | Illumina HiSeq 2500 | ERR1140785 |
| N0157 | L1.2.1 | Illumina HiSeq 2500 | ERR1140772 |
| N0157 | L1.2.1 | Illumina HiSeq 2500 | ERR1140791 |

**Supplementary Table 3.** Isolates used for the $\triangle$*hsdM* RNA-Seq analysis.

| ID | Lineage | Sequencer | NCBI ID |
|---|---|---|---|
| N1283-△hsdm | L4 | Illumina HiSeq 2500 | ERR2987806 |
| N1283-△hsdm | L4 | Illumina HiSeq 2500 | ERR2987808 |
| N1283-△hsdm | L4 | Illumina HiSeq 2500 | ERR2987810 |
| N1283 | L4 | Illumina HiSeq 2500 | ERR2987807 |
| N1283 | L4 | Illumina HiSeq 2500 | ERR2987809 |
| N1283 | L4 | Illumina HiSeq 2500 | ERR2987811 |

### Supplementary Results

#### 1. Precision and recall of Illumina data SV genotyping in complex and non-complex genomic regions

To further understand those regions where SVs can more accurately be genotyped using the *Mtb*-PRG and miniwalk approach, those that were found in the complex region masking scheme detailed in Marin *et al*[12] (~6% of the H37Rv genome length) were filtered out and precision and recall values were re-calculated. The precision of SVs in non-complex regions was of 0.81 and a recall of 0.32 in comparing against the SVIM-ASM truth set whereas the values were 0.89 and 0.42, respectively, against the *Mtb*-PRG truth set. Conversely, the precision and recall were also calculated for SVs in those complex regions resulting in 0.63 and 0.39, respectively, against the SVIM-ASM truth set and 0.69 and 0.39, respectively, against the *Mtb*-PRG truth set. These results show that genotyping SVs from short-read data can be highly accurate in non-complex regions and more caution should be taken in those more complex regions. However, it is important to note that the majority of SVs are found inside these complex regions.

#### 2. Extended DR analysis

For ethionamide (ETO), 5.3% of DR isolates had SVs in *ethA*, with the 1,655bp deletion being the most frequent. The other non-canonical gene (containing SVs) associated with ETO resistance was *Rv0221*, a gene previously associated with cycloserine (CS) resistance[1], though CS associations could not be tested due to low CS sample size.

For aminoglycosides, a 265bp deletion upstream *Rv2258c*, a possible transcriptional regulatory protein whose expression has been correlated with that of ribosomal protein genes[3], was associated with capreomycin (CAP) DR. SVs in *Rv2689c*, a 23S rRNA methyltransferase, were also associated with CAP DR, as well as amikacin (AMK) DR (Fig.S22). For streptomycin, a 220bp insertion upstream *rpoA*, an RNA polymerase, and downstream *rpsD*, a 30s ribosomal gene, was associated with DR. This insertion could potentially impact the expression of these ribosomal and transcriptional genes and counteract the drug's effect.

A gene associated with clofazimine (CFZ) resistance was *Rv1948*, the nucleotide diversity of which was shown to be increased in patients treated with bedaquiline (BDQ)[13]. An SV associated with delamanid (DLM) DR spanned *mycP2* and *eccE2*, both genes part of the ESX-2. The gene most significantly associated with DLM DR was *Rv0823c*, with 61bp deletions falling at the 5' start of the gene. However, these deletions are in a repetitive region where the resulting start codon is not affected by this deletion, though it could have an unknown effect on the gene's expression. This gene is a transcriptional regulator that forms part of the mycolic acids pathway (MAP) and is upregulated in response to MAP inhibitors[15]. DLM is a drug that targets the MAP pathway[8]; changes in *Rv0823c* gene expression could be associated to DLM DR. However, these deletions fall in a repetitive region where precision for genotyping small deletions is lower. The gene most significantly associated with bedaquiline (BDQ) resistance was *Rv1998c*, a probable phosphonomutase.

Moreover, we found SVs in genes already associated with resistance which did not reach significance due to a lack of power, for example a 438bp deletion found in two bedaquiline-resistant isolates spanning the start codon of the *mmpR5* gene or two para-aminosalicylic acid-resistant isolates with 2 different deletions spanning the *thyA* gene. A 7,945bp deletion spanning *katG* was also observed in 11 isoniazid-resistant isolates.

Most DR-associated variants were filtered out for having a minimum allele frequency (MAF) $< 0.01$. However, we found an interesting 102bp deletion in *hsdM* with MAF ~0.005 highly associated with INH DR (Fig.S20a). HsdM is a methyltransferase which has been previously associated with isoniazid resistance in knock out studies[4]. This deletion, though it does not cause a premature stop codon, affects the protein's N6 adenine-specific DNA methyltransferase N-terminal domain (UniProt ID: A0A655I246), as well as the rest of the protein structure (Fig.S20b), possibly rendering it non-functional. To study the effect of a loss of function of this protein in drug resistance, we analysed the transcriptional profile of an isolate with *hsdM* knocked out(*hsdM*-DEL) against the same isolate with complete *hsdM*[2]. Though not significant (adjusted $p$-value=0.056), *inhA* was enriched with a fold change of 0.25 in *hsdM*-DEL (Fig.S20c). Overexpression of *inhA* has been previously attributed to INH resistance[10]. These results point towards possibly low-level INH DR. Interestingly, *ethA* was significantly depleted (adjusted $p$-value=0.034) in *hsdM*-DEL, a gene whose deletion we found to be associated with ETO resistance, though the *hsdM* deletion was predominantly found in ETO-susceptible isolates (Fig.S20d).

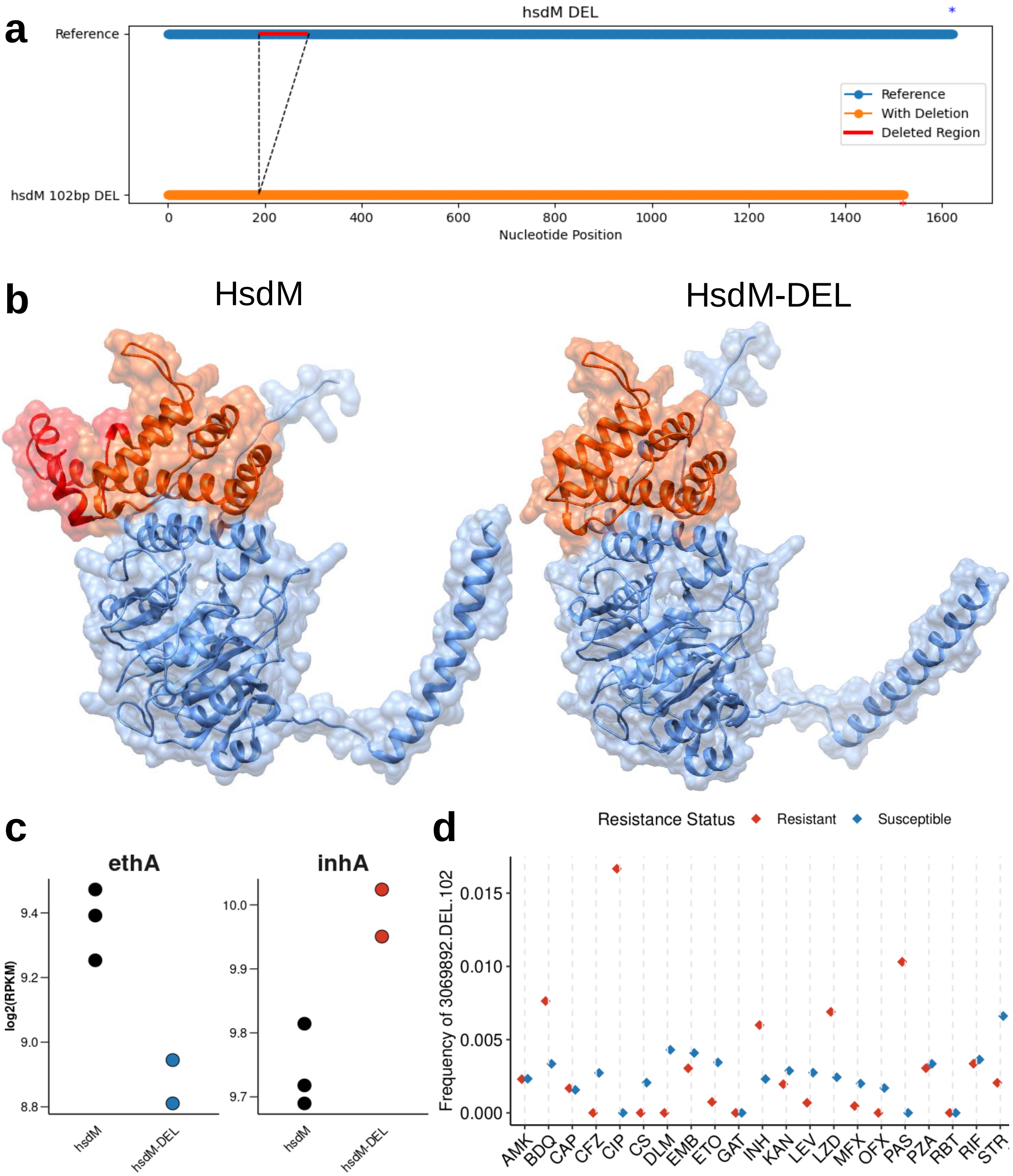


**Supplementary Figure 20**. Impact of the 102bp *hsdM* deletion associated with INH resistance. **a** 102bp deleted sequence from the reference genome *hsdM*. Asterisks represent stop codons. **b** Protein structure of *hsdM* in the reference genome (left) and of *hsdM* with the deletion (right). The orange area represents the N6 adenine-specific DNA methyltransferase N-terminal (UniProt: A0A9P1L9Y1). The red area represents the deletion. Both structures were predicted using Alphafold (v2.3.2). **c** Dot plots showing expression patterns for *ethA* and *inhA* in a *hsdM*-knock out (*hsdM-DEL*) against the same isolate without the knock out. In red are genes enriched and in blue are genes depleted in the knock out. **d** Frequency of the 102bp *hsdM* deletion across resistant and susceptible isolates to 21 different drugs. Acronyms are defined in Fig.5.

### Supplementary Methods

#### 1. Steps to genotype SVs using minigraph and miniwalk

To genotype SVs using minigraph-miniwalk, 6 steps should be taken:

1. Create an assembly (i.e. with Flye [9] if using long-read data or with Shovill [16] if using short-read data).
2. Find the paths the assembly goes through using `minigraph -cxasm --call graph.gfa assembly.fasta > paths.bed`.
3. The bed-like file resulting from the previous mapping is then merged into a bed file with the reference paths using minigraph's ”merge” function and posteriorly transformed into a VCF file using the ”merge2vcf” function.
4. Use `mod` module from miniwalk. This code takes the VCF output from the previous step to obtain a VCF with specific SVs and their positions.
5. Use `ref` module from miniwalk. This code takes the output VCF from `mod` and outputs a new VCF file with the purpose of ordering the SVs by their position and grouping those in close proximity into a single SV line.
6. Use `ins2dup` module from miniwalk. This code reads a VCF file and goes through all the called INS SVs and determines whether they are a DUP instead.

#### 2. Miniwalk code detailed description

- `mod`
  This code takes the output from the ”merge2vcf” minigraph function to obtain a VCF with specific SVs and their positions. It requires 4 inputs: `-b/--bed` (the bed file output from `minigraph -xasm --call`), `-v/--vcf` (the VCF file output from `merge2VCF`), `-g/--gfa` (the gfa pangenome file), `-o/--output` (the new VCF file to store the SVs' information) and an optional `-na` flag. The code starts by looking if there is a ”.” in the bed file, indicating that that specific bubble has not been resolved, and if that is the case then that specific SV region will be marked with NAs (in case the `-na` flag has been selected). If that is not the case, we continue on to classify each SV. Those bubbles that hold an SV are marked in the input VCF as ”<X:1>”, therefore those will be the ones that go on to the final VCF. A bubble that holds a ”*” means that there is only one SV; depending on whether the asterisk is on the reference path or on the sample path, the SV will be classified as ”INS” or ”DEL”, respectively. The SV length will be determined by the sum of each node's length in the opposite path from the asterisk. In the case a bubble holds two different paths (as opposed to a path and an asterisk), it means that there may be more than one SV. Each path is separated into the different nodes that make it up, and subsequently compared to the opposing path's nodes to determine which nodes are unique to the path, meaning that an SV lies there. Nodes unique to each path are further grouped into those that are sequential to each other. Each group of nodes sequential to each other make up a candidate SV as a whole; those found only in the reference path are considered as deletions while those in the sample path are considered as insertions, initially. After grouping the nodes, each group is looked at in more detail to determine the size of the SV as well as the type of SV, as inversions can be incorrectly classified as insertions. To look for inversions, there are two ways to determine them; if on the reference path there is a node which also appears on the sample path, but with the opposite orientation (i.e. holds a ”<”), there is an inversion. The other, slightly more complex method to find an inversion is to go through every node inside the current SV grouping and map its reverse complement to the nodes in the opposite path; if the mapping accuracy is of 95% with a length difference error margin of 5%, the node is classified as an inversion and the current SV is separated into two SVs if the node is found in the middle of the SV grouping. If no inversion is detected, the SV is classified as an insertion. Insertion length and specific sequence is determined by sequentially adding the nodes in their predetermined order. Sequences of nodes that present a ”<” are transformed into their complementary reverse sequence and subsequently added to the rest of the sequence. The same process, save from the inversion determination, is followed with the reference path to classify SVs as deletions. The flowchart of this script is described in Fig.S21.

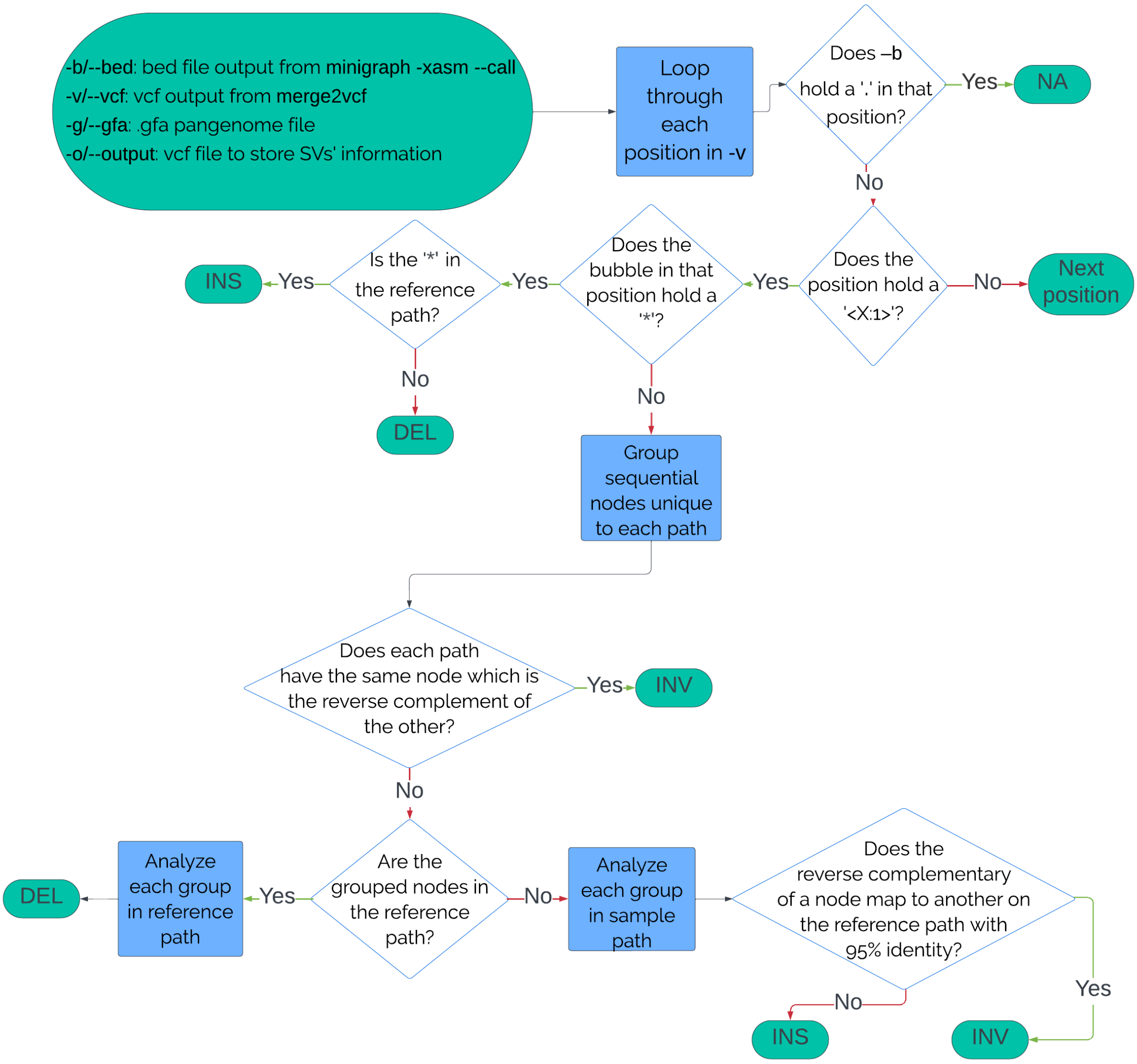


**Supplementary Figure 21**. Flowchart of the `mod` miniwalk code.

- `ref`
  This code takes the output VCF from `mod` and outputs a new VCF file with the purpose of ordering the SVs by their position and grouping those in close proximity into a single SV line. The code starts by ordering SVs by their specific position. If an insertion's length covers one or more deletions in the same region of the genome, those deletions are grouped together and will appear in the same line as the insertion in the output VCF; the SV will be classified as an insertion if the insertion remains larger than the grouped or individual deletions. As certain insertions or deletions do not completely overlap each other, 150bp error overlap margin is allowed. An exception is created with two insertions, as they will only be clustered if they are ≤250bp in proximity of each other, not taking their size into account. If the difference in size of the reference sequence against the sample sequence is smaller than 50bp, though the sequences may be different in content, the SV is not added to the final VCF as the SV is not large enough. The whole process works vice versa, i.e. in case a deletion encompasses one or more insertions. In those cases where the ref and sample SVs have a sequence that is highly similar at the beginning of the larger sequence, the sequences will be mapped locally using the pairwise alignment tool from Biopython to determine whether the exact breakpoint should be further down the genome and if so, modify it. In the case that the sequences map at the end of the larger sequence, the SV breakpoint will not need to be changed. The flowchart for this script is described in Fig.S22.

- `ins2dup`
  This code reads a VCF file and goes through all the called INS SVs and determines whether they are a DUP instead. For each insertion, an extended INS allele is created by extracting reference sequence from the H37Rv genome immediately upstream (2x variant size) and downstream (2x variant size) of the variant site, then concatenating these in appropriate order with the consensus insertion sequence. Each extended allele is then scanned for tandem repeats using the `repeat-match -t` function from MUMmer [11]. Those INS that have tandem repeats of at least half the size of the INS are considered as DUP and are changed accordingly in the VCF file.

Another Python code was created to do the benchmarking, as there is no consensus tool for benchmarking SVs, moreover the SVs output by the graph genotyping approach are not compatible with the few current tools available. The code is explained as follows:

- `bench`
  This code outputs the precision and recall when comparing variants from a truth set (SVIM-ASM) and a call set (minigraph). The inputs required are `-v/--vcf` (truth set), `-c/--call` (call set), `-b/--bed` (bed file to identify any NAs), `-r/--repeat` (tandem repeat regions) and `-e/--reference` (reference used to get position coordinates). Tandem repeat regions were determined by using nucmer's repeat-match command and keeping those repeats which were overlapping each other or maximum 100bp from each other [11]. The code firstly goes through each SV in the call set individually, comparing it to those SVs in the truth set. A SV is classified as a true positive (TP) when it is overlapping the truth set SV by at least 25% and is the same SV type [5, 18]. If an SV does not fulfill the aforementioned requirements but is found inside a tandem repeat (TR) region, the 50 contiguous base pairs will be extracted along with the SV for both the call and truth set found inside the same TR region and mapped against each other; we will classify the SV as a true positive if the sequence overlap is of at least 50%. If an SV in the call set spans two truth set SVs (or vice versa) which are contiguous to each other by < 200bp and the sum of their sizes is inside the call set SV's 50% error margin, the SV is classified as a TP. Those call set SVs that do not fulfill any of the previous requirements are classified as false positives (FP). Lastly, each SV in the truth set is analysed to determine whether any SVs were missed, classifying them as false negatives (FN). SVs that were classified as "incomplete_inversions" are not added as FN. The flowchart for this script is shown in Fig.S23.

To validate that our code was working, 5 different synthetic datasets were created; a SVIM-ASM truth VCF, a *Mtb*-PRG truth VCF, a long-read *Mtb*-PRG VCF, a short-read *Mtb*-PRG VCF and a manta VCF. We added different types of SVs (INSs, DELs and INVs) as well as different scenarios; an SV that was exactly like the gold standard, a false positive SV, a false negative SV, a called SV that did not overlap with the standard SV but was in the same tandom duplication with the exact same sequence and, therefore, should be a true positive, a called SV that overlaps 2 standarad SVs or vice versa, a called SV that entirely spans a standard SV and overlaps ≥25% and a called SV that is spanned by a

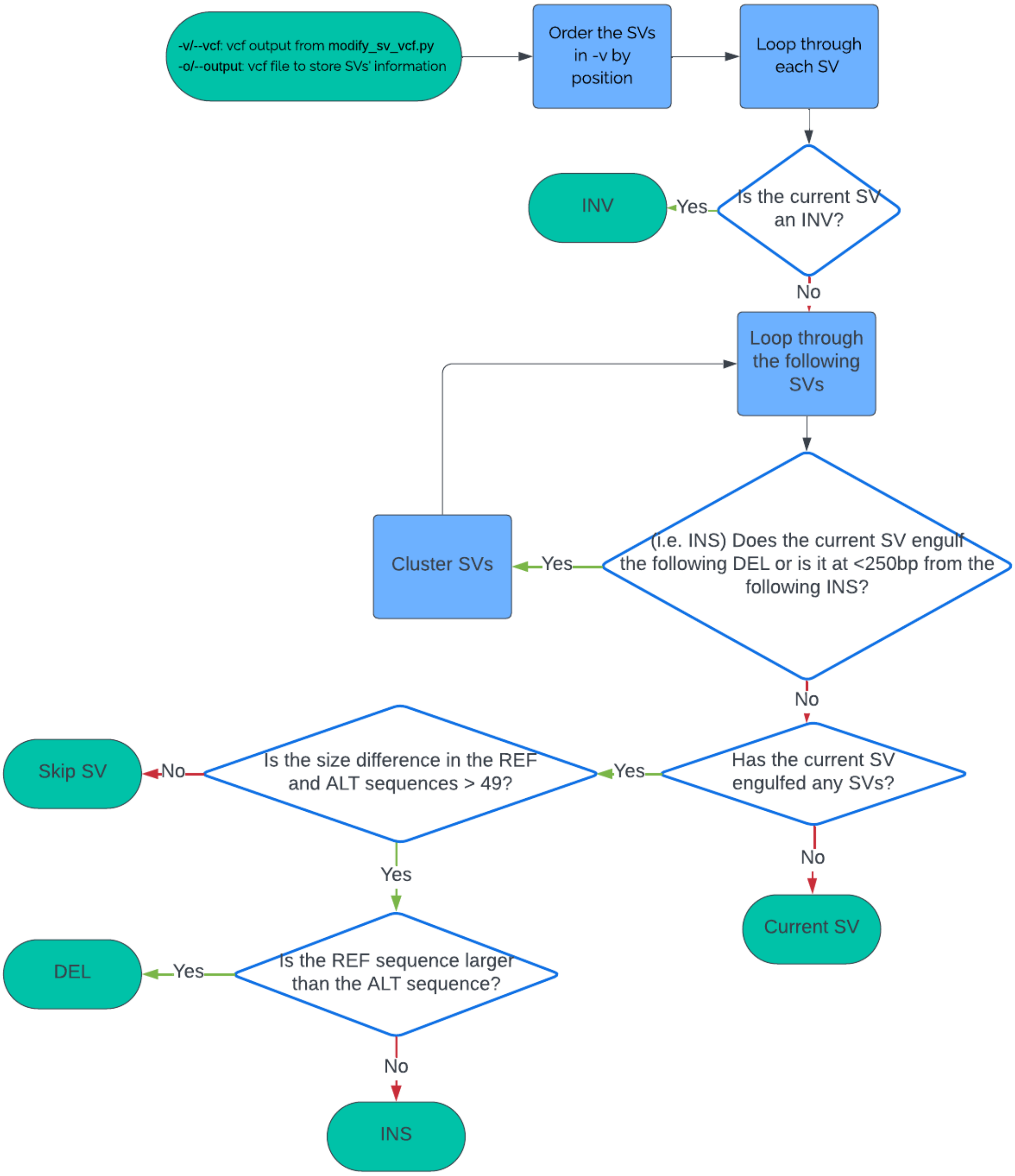


**Supplementary Figure 22**. Flowchart of the `ref` miniwalk code.

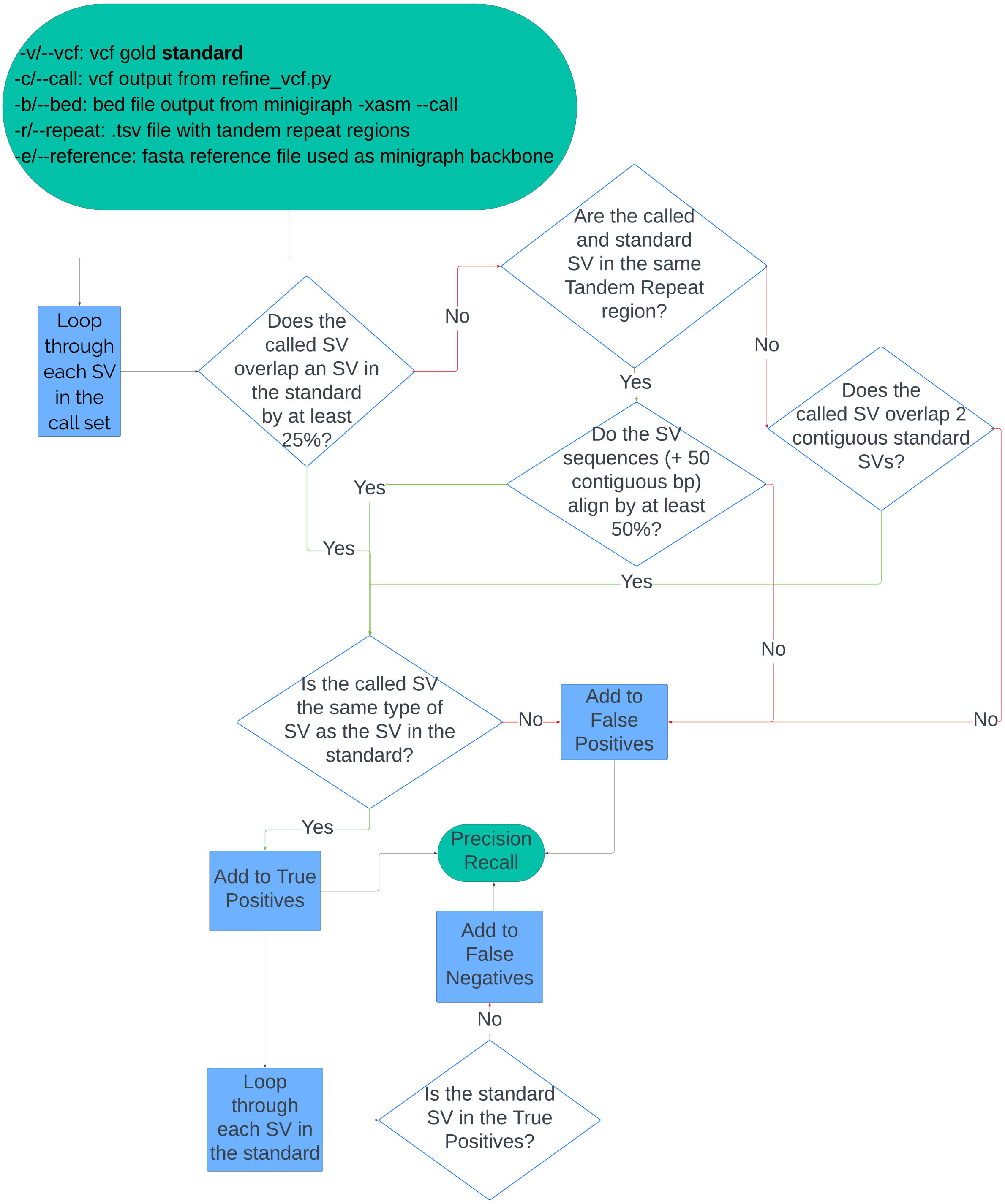


**Supplementary Figure 23**. Flowchart of the `bench` code.

standard SV but overlaps <25%. After finding that the code output the expected precision and recall values, the code was used for the benchmarking.

#### 3. HsdM protein visualisation

Chimera (v1.18)[14] was used to visualize the protein structure using predicted protein structures with and without the *hsdM* deletion from Alphafold (v2.3.2)[6] predictions.

#### 4. *hsdM* knock out transcriptome analysis

RNA-Seq analyses were carried out as explained in "L1.2.1 transcriptome analysis". Data was downloaded from[2] (Table S3). A replicate whose expression did not correspond with the other two was filtered out (Fig.S24).

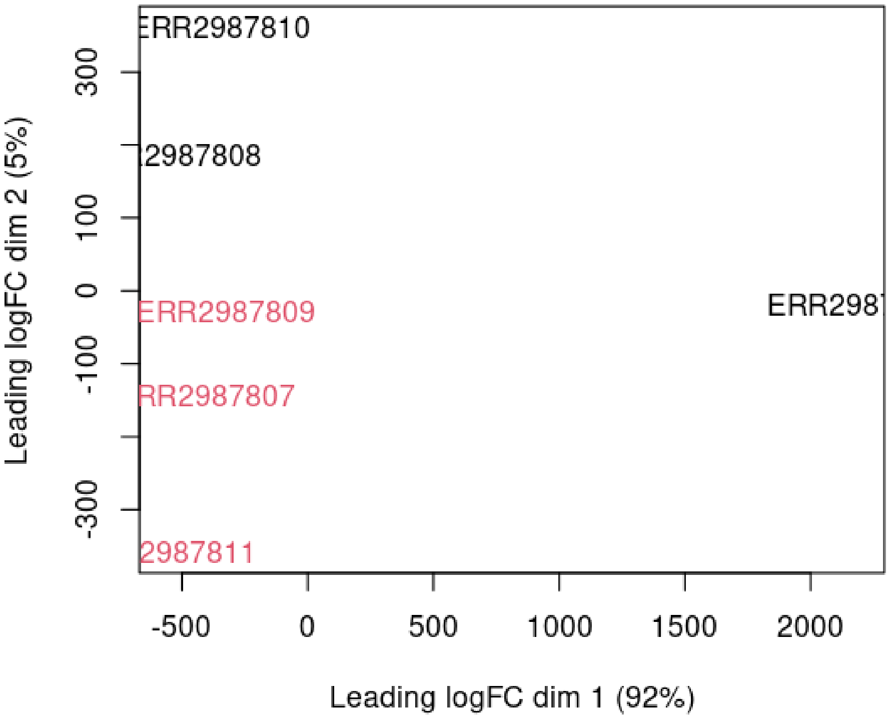


**Supplementary Figure 24.** A multidimensional scaling (MDS) plot showing the 3 replicates of two isolates (black and red). An MDS plot is a visualisation of a principle components analysis (PCA), which determines the greatest sources of variation in the data.
